# Cell-type specific concentration regulation of the basal transcription factor TFIIH

**DOI:** 10.1101/613281

**Authors:** Lise-Marie Donnio, Catherine Miquel, Wim Vermeulen, Giuseppina Giglia-Mari, Pierre-Olivier Mari

## Abstract

The basal transcription/repair factor TFIIH is a ten sub-unit complex essential for RNA polymerase II (RNAP2) transcription initiation and DNA repair. In both these processes TFIIH acts as a DNA helix opener (by the enzymatic activity of the XPB and XPD helicases), required for promoter escape of RNAP2 in transcription initiation, and to set the stage for strand incision within the Nucleotide Excision Repair (NER) pathway. We generated a knock-in mouse model that endogenously expresses a fluorescent version of XPB. Here we demonstrate, via confocal imaging of ex vivo tissues and cells derived from this mouse model, that TFIIH steady state levels are tightly regulated at the single cell level, thus keeping nuclear TFIIH concentrations remarkably constant in a cell type dependent manner. Moreover, we show that individual cellular TFIIH levels are proportional to the speed of mRNA production, hence to a cell’s transcriptional activity, which we can correlate to proliferation status. Importantly, cancer tissue presents a higher TFIIH than normal healthy tissues. Taken together, these results show that TFIIH cellular concentration might be used as a *bona-fide* marker of transcriptional activity and proliferation.

**Significance:** Using a mouse model expressing a fluorescently version of TFIIH, we showed that TFIIH concentration is tightly controlled and that this concentration is proportional to the cellular transcriptional activity and proliferation capacity.

## Introduction

The integrity of the genome is continuously challenged by a variety of genotoxic agents, causing DNA lesions, which interfere with transcription (inducing premature cell death and ageing) and DNA replication (inducing mutations leading to cancer). To prevent the deleterious consequences of genomic insults, genome surveillance systems including DNA repair processes protect the genomic information. Nucleotide excision repair (NER) is one of the most important DNA repair systems and removes helix-distorting DNA-adducts in a complex multi-step reaction. Disruption of NER in response to genotoxic injuries results in the rare genetic disorders ranging from the cancer-prone xeroderma pigmentosum (XP) phenotype to premature ageing Cockayne syndrome (CS) and Trichothiodystrophy (TTD) (1).

In the NER pathway, the transcription factor TFIIH unwinds DNA and recruits other factors required to remove the damaged oligonucleotide. TFIIH is a 10 subunit protein complex (2) essential for RNA polymerase II (RNAP2) transcription initiation (3), RNA polymerase I (RNAP1) transcription (4) and NER (5). The ten subunits cluster in two sub-complexes: the core complex composed of 7 tightly bound proteins (XPB, XPD, p62, p52, p44, p34 and TTDA) and the CAK (CDK-activating kinase) complex composed of the remaining 3 components (CDK7, Cyclin H and MAT1), which are more loosely bound to the core via their interaction with the XPD protein.

Mutations in the TFIIH subunits XPB, XPD and TTDA have been associated with the three diseases: Xeroderma Pigmentosum (XP [MIM 278700-780]), Cockayne Syndrome (CS [MIM 214150]) and Trichothiodystrophy (TTD [MIM 601675]), all characterized by deficiency in DNA repair (6). However, despite a similar DNA repair defect, while XP patients are highly cancer prone, CS and TTD phenotypes are not associated with cancer predisposition. This discrepancy cannot simply be explained by a DNA repair defect and could be the result of the affected TFIIH transcriptional functions (7). Besides the common DNA repair defect, a characteristic of TTD patients mutated in TFIIH subunits is the low cellular TFIIH steady state levels (2), in contrast with the cancer-prone XP patients that have normal TFIIH levels.

Importantly, it is nowadays acquired (mostly in cancer related research) that a bidirectional connection exists between cellular metabolism and gene transcription and that both are probably tightly coordinated (8). The increased cellular metabolism needed for chronic unscheduled proliferation in cancer cells induces and depends on an increased transcriptional activity and in many cancers a dysregulation of the RNAP2 and RNAP1 transcription have been observed (9). The importance of transcriptional activity during carcinogenesis is best highlighted by the fact that many molecules interfering with transcription have been used as chemotherapeutic drugs (10) and particularly two chemotherapeutic drugs, triptolide and spironolactone, interfere with TFIIH activity (11, 12).

On the basis of these observations, it has been hypothesised that reduced TFIIH levels could play a protective role by preventing chronic proliferation: a mandatory requirement during carcinogenesis (2). In order to verify this hypothesis, a first step would be to demonstrate that TFIIH cellular concentration is indeed linked to transcriptional activity and cellular metabolism/proliferation. To verify this hypothesis, we analysed TFIIH steady state levels in a mouse model system in which the yellow fluorescent protein (YFP) was targeted to the last exon of the *XPB* gene (encoding for one of two helicase that constitute the core TFIIH complex, hereafter XPB-YFP) (13). Homozygous XPB-YFP knock-in mice (Xpb^y/y^) were born healthy, fertile and DNA repair efficient (13) and no features of premature ageing or increased spontaneous carcinogenesis were observed (13).

Our results show that TFIIH cellular concentration is strictly regulated in distinct cell types within the organism. Importantly, we could show that TFIIH steady state correlates with transcriptional activity and cellular proliferation and that cancerous tissues show increased TFIIH levels.

## Results

### Single cell quantification of TFIIH in organs from the Xpb^y/y^ mouse model

We used a mouse model system in which the yellow fluorescent protein (YFP) was targeted to the last exon of the *XPB* gene (encoding for one of two helicase that constitute the core TFIIH complex, hereafter XPB-YFP) (13). Homozygous XPB-YFP knock-in mice (Xpb^y/y^) were born healthy, fertile and DNA repair efficient (13) and no features of premature ageing or increased spontaneous carcinogenesis were observed (13).

In order to map TFIIH levels in a living organism, direct measurements of the XPB-YFP fluorescence in both cryosections of organs and living organotypic tissue slices were undertaken, on our Xpb^y/y^ mouse model, via 3D confocal microscopy. The choice of using organs and cells derived from the Xpb^y/y^ mouse model and to directly measure the YFP fluorescent signal was made after having considered that as a core sub-unit of TFIIH, XPB is considered to exist in the nucleus only as part of TFIIH and this holds true for the fluorescently tagged XPB (14). Importantly, the presence of the YFP fluorescent tag in fusion with XPB within TFIIH does not affect TFIIH steady state as shown in Figure S1, demonstrating that our approach and model are both solid and that results can be extrapolated to the untagged TFIIH complexes *in vivo*. Additionally, the direct measure of the YFP fluorescence circumvents possible measurement alterations caused by antibody amplification of signals.

The YFP derived signal was detectable in cryosections of all the tissues we tested (brain, intestine, kidney, liver, spleen, skin and testis). Initial analysis showed that different cell types could have very different XPB-YFP expression levels. For example, in the brain cortex, glial cells present just 20% of the YFP signal measured in adjacent neurons (Fig. 1A and 1B). The much lower transcriptional activity of glial cells, under unchallenged conditions, compared to neurons (15), suggests that the amount of TFIIH in cells could reflect specific neuronal transcriptional activity.

**Figure 1.**
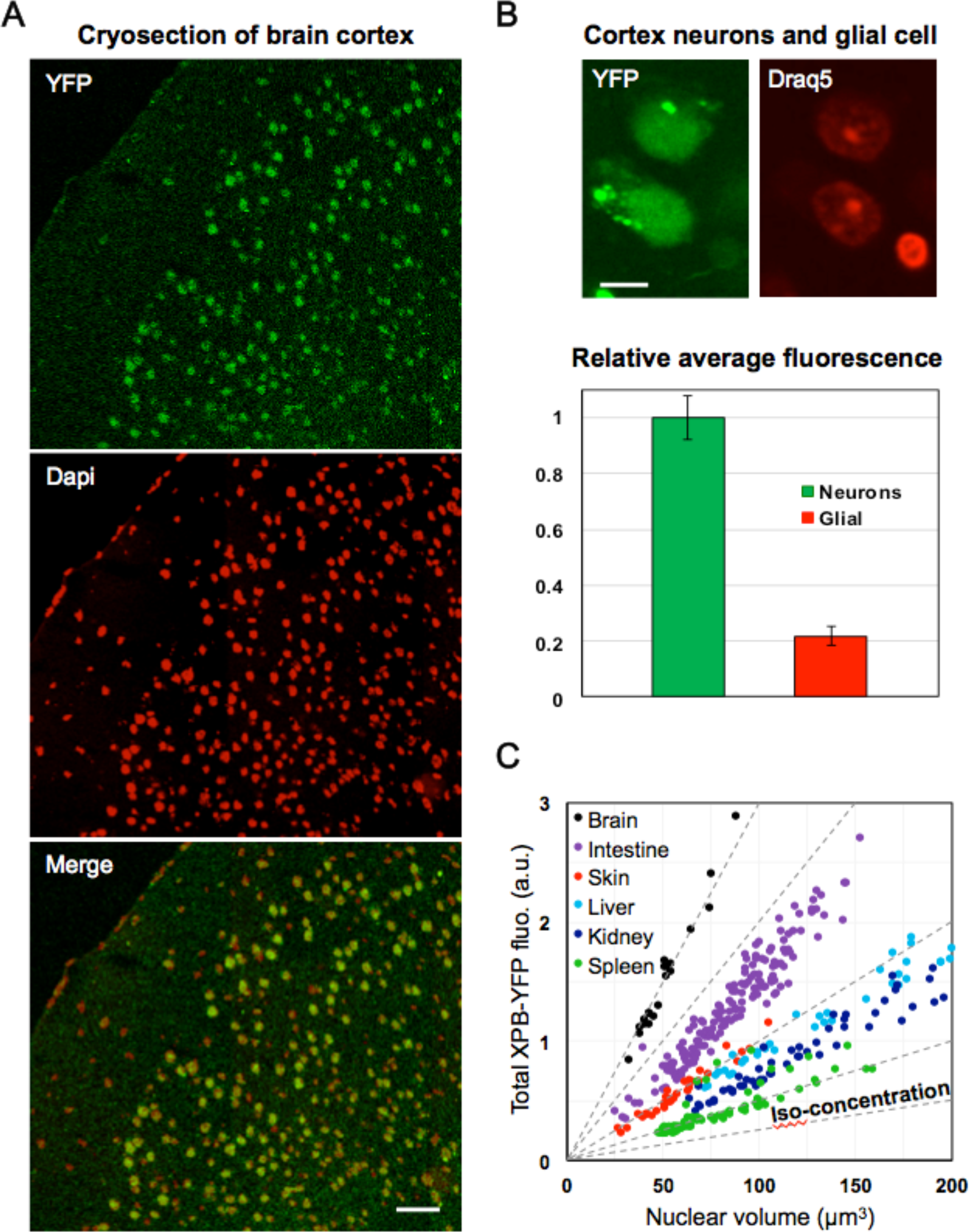
Differential XPB steady state levels in the Xpb^y/y^ knock-in mouse model. **A.** Low magnification confocal image mosaic of a cortex cryosection (Xpb^y/y^ mouse), showing the YFP signal in green and Dapi in red to enhance contrast in the merged image. Individual cells are singled out by the strictly nuclear XPB-YFP and/or Dapi signals. Background and auto-fluorescence limits the YFP detection of weakly expressing cells that appear orange/red in the merged image. **B.** Neurons and a Glial cell imaged within a living organotipic slice of cortex (Xpb^y/y^ mouse). The Glial cell is easily distinguished from the neurons by its smaller nucleus size and condensed chromatin (15) dyed with the supravital DNA marker Draq5 (Biostatus, UK). The measured average nuclear YFP signal in both cell types (N=24) is plotted in the accompanying graph, which shows a 5-fold difference in the TFIIH concentration between these two cell types. **C.** Plot of the total XPB-YFP signal per cell versus the reconstructed volume occupied by the YFP signal of each analysed cell. In each of these tissues, cell to cell variations of the total quantity of XPB-YFP and of the nuclear volume cancel out in a non-trivial fashion, resulting in cells with the same nuclear XPB-YFP concentration. Radial dotted lines represent constant XPB-YFP concentrations.

To reliably assess whether or not TFIIH levels correlate with transcriptional activity, we first needed to develop a quantitative approach that allows more accurate measurements of the *total* amount of XPB-YFP per cell nucleus. To achieve this, we used a confocal laser-scanning microscope to acquire 3D image stacks of tissues and cultured cells derived from the Xpb^y/y^ mouse model. Typically, *ex vivo* tissue slices were maintained metabolically active during image acquisition under the microscope (13). The resulting 3D image sets were processed via quantitative deconvolution in order to spatially reassigns “out-of-focus fluorescence” to the correct voxel coordinates. The total amount of TFIIH per cell was then estimated by integrating the XPB-YFP fluorescence signal within each fully imaged nucleus (cf. Fig. S2 and Methods).

The results we obtained for several tissues (brain, intestine, kidney, liver, skin and spleen) are shown in Figure 1C, with the total fluorescence per nucleus plotted (in arbitrary units) versus the reconstructed volume occupied by the YFP signal of each analysed cell (in cubic micrometers). By displaying our data in this manner, a linear relation between the total amount of TFIIH per cell and the nuclear volume is revealed. The existence of such a strong correlation is non-trivial and indicates that TFIIH steady state levels are tightly regulated at the individual cell level *i.e.* two cells of the same type but with different nuclear volumes will have different TFIIH expression levels in order to compensate for nuclear volume differences, thus maintaining a constant TFIIH concentration (concentration iso-contours appear in Fig. 1C as radial dashed lines). The relative average TFIIH concentrations of cells in different tissues are easily discriminated (Fig. 1C). In decreasing order, we find: brain (neurons), intestine (epithelial), skin (epidermal keratinocytes), liver (hepatocytes), kidney and spleen. This level of regulation is not observed in SV40-immortalized human fibroblasts, stably expressing XPB-GFP under an exogenous promoter (14), hence this unique “constant concentration” signature has yet to be observed outside of the Xpb^y/y^ mouse model context.

### TFIIH levels correlate with transcriptional activity

A clear example suggesting a correlation between transcriptional activity and TFIIH levels was found when inspecting sections of testis, in particular the seminiferous tubules in which male germ cell development proceeds in a peripheral-to-central fashion. Indeed, during this process, chromatin in male germ cells is progressively condensed, concomitantly genes are silenced by promoter methylation and gradually basal transcription is reduced to undetectable levels (16–18). Immunohistochemistry of the seminiferous tubules of an Xpb^y/y^ mouse showed that TFIIH concentration follows the peripheral-to-central reduction of transcriptional activity (Fig. 2A), with mature sperm cells, observed in the lumen of the tubule, not showing any detectable TFIIH signal.

**Figure 2.**
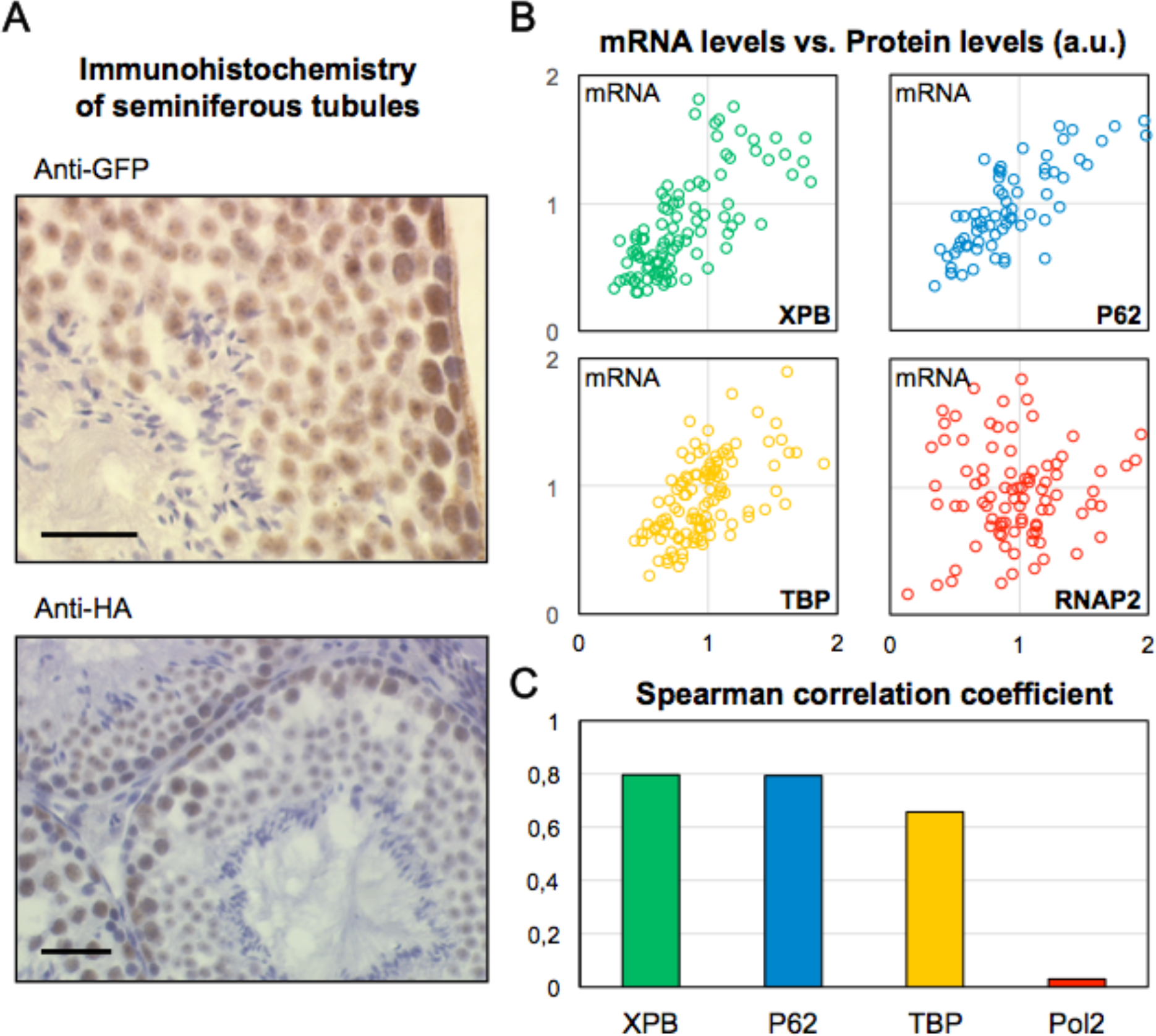
Transcriptional activity correlates with TFIIH levels. **A.** Sections of testis (Xpb^y/y^ mouse) showing seminiferous tubules were stained with either anti-GFP or anti-HA (DNA content as counter staining). In both images, TFIIH concentration (via XPB) follows the known peripheral-to-central reduction of transcriptional activity. Mature sperm cells in the lumen of the tubules only appear in counter staining (no detectable TFIIH). Scale bars 50 µm. **B.** Plots of the total amount of mRNA versus the total quantity of XPB (green), p62 (blue), TBP (yellow) and RNAP2 (red), per nucleus (in arbitrary units) for Mouse Dermal Fibroblasts. N= 111; 72; 119 and 94 respectively. **C.** Plot of the corresponding Spearman correlation coefficient between mRNA and XPB, p62, TBP and RNAP2 levels. RNAP2 and mRNA levels do not appear correlated.

So far, the evidence that TFIIH quantity is related to transcriptional activity is indirect and based on observations conducted on different murine tissues, known to present different cell types with distinct transcriptional programs.

To test whether transcriptional activity does indeed correlate with TFIIH levels, we measured mRNA production by pulse-labelled newly transcribed RNA in cultured cells by Bromo-Uridine or 5-ethynyl Uridine (EU) incorporation. The incorporation time was kept under 30 minutes in order to minimize cytoplasmic export of the produced mRNA, hence dilution throughout the cell body. We estimated concomitantly EU and TFIIH amounts in cultured Xpb^y/y^ Mouse Dermal Fibroblasts (MDF) by immunostaining of TFIIH subunits and click-it reaction of the EU-labelled mRNA (Fig. 2B, green) with our validated 3D quantitative imaging approach. In figure 2B (green dots), a linear correlation can be observed between the measured amount of mRNA produced and XPB steady-state levels. We could also measure the same correlation when cells were immuno-stained with an antibody against p62, another TFIIH core subunit (Fig 2B, blue dots), thus demonstrating that the total amount of cellular TFIIH is proportional to the cellular transcriptional activity. Interestingly, we could also measure the correlation between mRNA production and the basal transcription factor TBP (Tata-Binding Protein) (19) that is part of the TFIID complex, which also participate in RNAP2 transcription initiation, and we could also measure a correlation (although less proportional that the one observed for TFIIH) between the total amount of TBP and the transcriptional activity (Fig 2B, yellow dots). In contrast, no correlation between transcriptional activity and amount of RNAP2 was found (fig 2B, red dots), showing that not all transcription factors follow the same strict correlation rules and that TFIIH and TBP might be good indicators/markers of the cellular transcriptional activity. The Spearman correlation coefficient of the four data set is recapitulated in Fig 2C.

### TFIIH levels correlate with proliferation

Since TFIIH can provide a measure of transcriptional activity, we also wanted to determine if TFIIH can also be used as a marker for proliferation. To establish this, we modulated the proliferation status of cultured cells then measured changes in mRNA production and TFIIH steady-state levels. Modulation of proliferation was obtained in cultured primary chondrocytes (neither transformed nor immortalized) kept in low oxygen (3%) culture conditions. Serum starvation of confluent chondrocytes cultures will strongly stop proliferation, while dilution of these cultures in presence of a high level of serum will rapidly induce proliferation. Our measures of mRNA production and total TFIIH amounts show that in murine chondrocytes (Fig. 3A) isolated from the Xpb^y/y^ mouse model, arrested cells present a low basal transcriptional activity and correspondingly low TFIIH amounts. On the contrary, the general mRNA production and the total cellular TFIIH amounts are higher in proliferative chondrocytes (Fig 3A). We verified the results by Western Blot of the subunits XPB, p62, CDK7 and Cyclin H in both culture conditions (Serum-starved and Proliferation) (Fig 3B). Because different TFIIH levels can be the result of an increased transcriptional activity hence an increased protein production, we loaded the same number of cells per well (500.000). A clear increase of core and CAK subunits of TFIIH have been observed in the proliferative cells compared to the serum-starved. Indeed, when the TFIIH level of serum-starved cells was normalised to 1, an increase of 50 to 80 % was measured in proliferative cells (Fig 3C).

**Figure 3.**
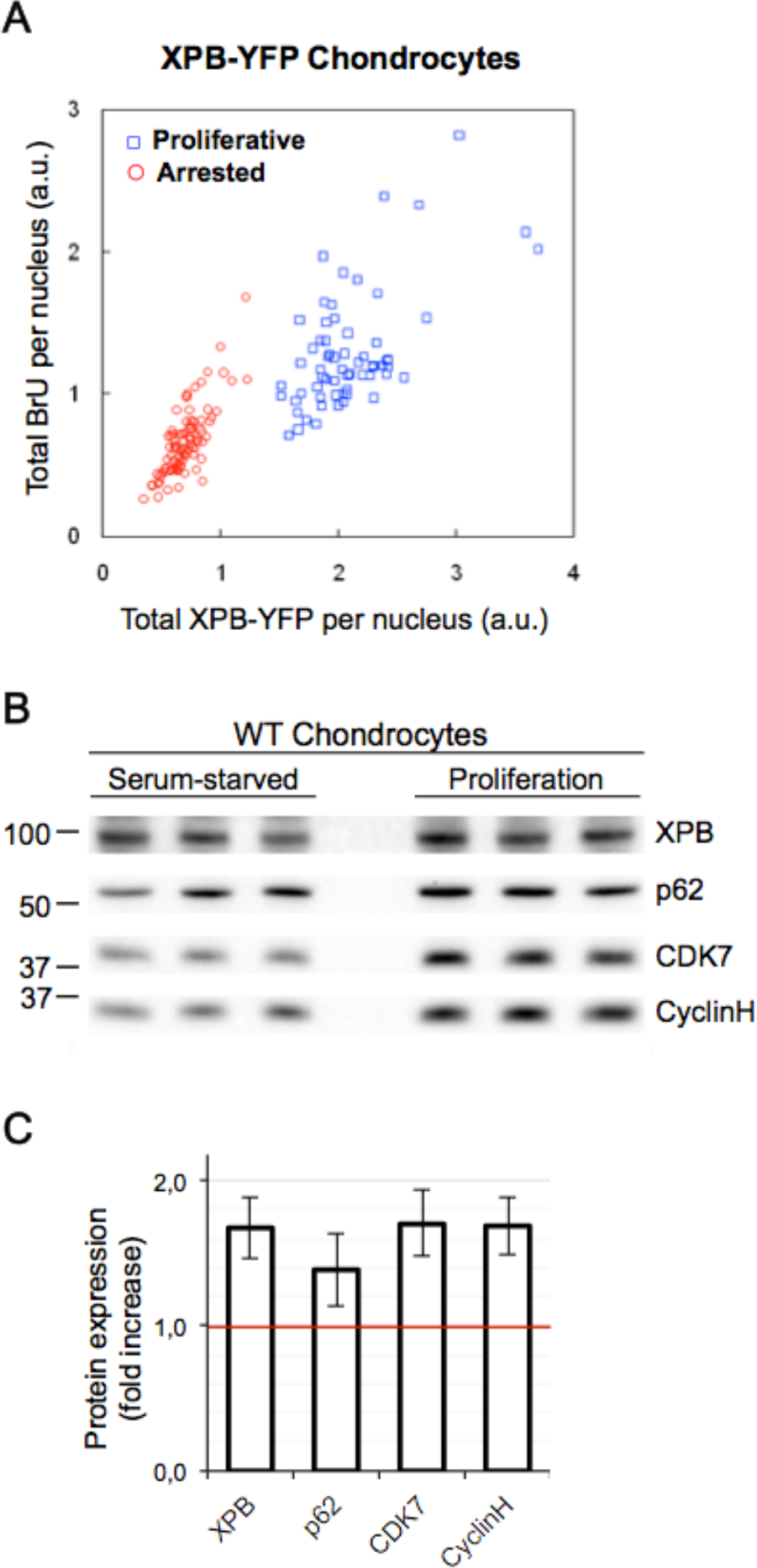
Proliferation correlate with TFIIH levels. **A.** Plot of the total amount of BrU incorporated mRNA versus the total quantity of XPB-YFP (per nucleus, in arbitrary units) in quiescent (red circles) and proliferating (blue squares) XBP-YFP chondrocytes. The increase in XPB levels concomitant with the increased BrU levels (associated to the transcriptional boost in proliferating cells) is clearly visible. **B** Western blot showing the expression of TFIIH subunits in whole cell extracts of Chondrocytes in serum-starved versus proliferative state. The same quantity of cells was used for extraction and loading. **C** Graph combining at least three different Western blots showing the fold increase of TFIIH subunit expression levels in proliferating cells normalized to serum-starved cells.

### TFIIH levels in colorectal cancer

In order to investigate whether TFIIH levels are also increased in cancer tissues, we stained a tissue section of a human colorectal cancer including adjacent normal colon tissue with an antibody against the XPB subunit of TFIIH and the proliferation marker Ki67 (Fig. 4). The expression patterns in both tumor and non-tumor tissues were evaluated by the pathologist of this study. We found that XPB and Ki67 were concomitantly highly overexpressed in primary colon cancer tissue (ROI 1, Fig. 4A and 4B) in strong contrast to adjacent normal colonic tissue (ROI 2, Fig. 4A and 4C) where both XPB and Ki67 were only minimally detected. Interestingly, in normal adjacent colon tissue, we observed a few cells with high levels of XPB correlated with Ki67 expression (Fig. 4C), in the deeper regions of colonic crypts, where proliferating cells are expected. A consecutive serial section was also stained with hematoxylin to evaluate the morphology of the tissue (Fig. 4D, 4E).

**Figure 4.**
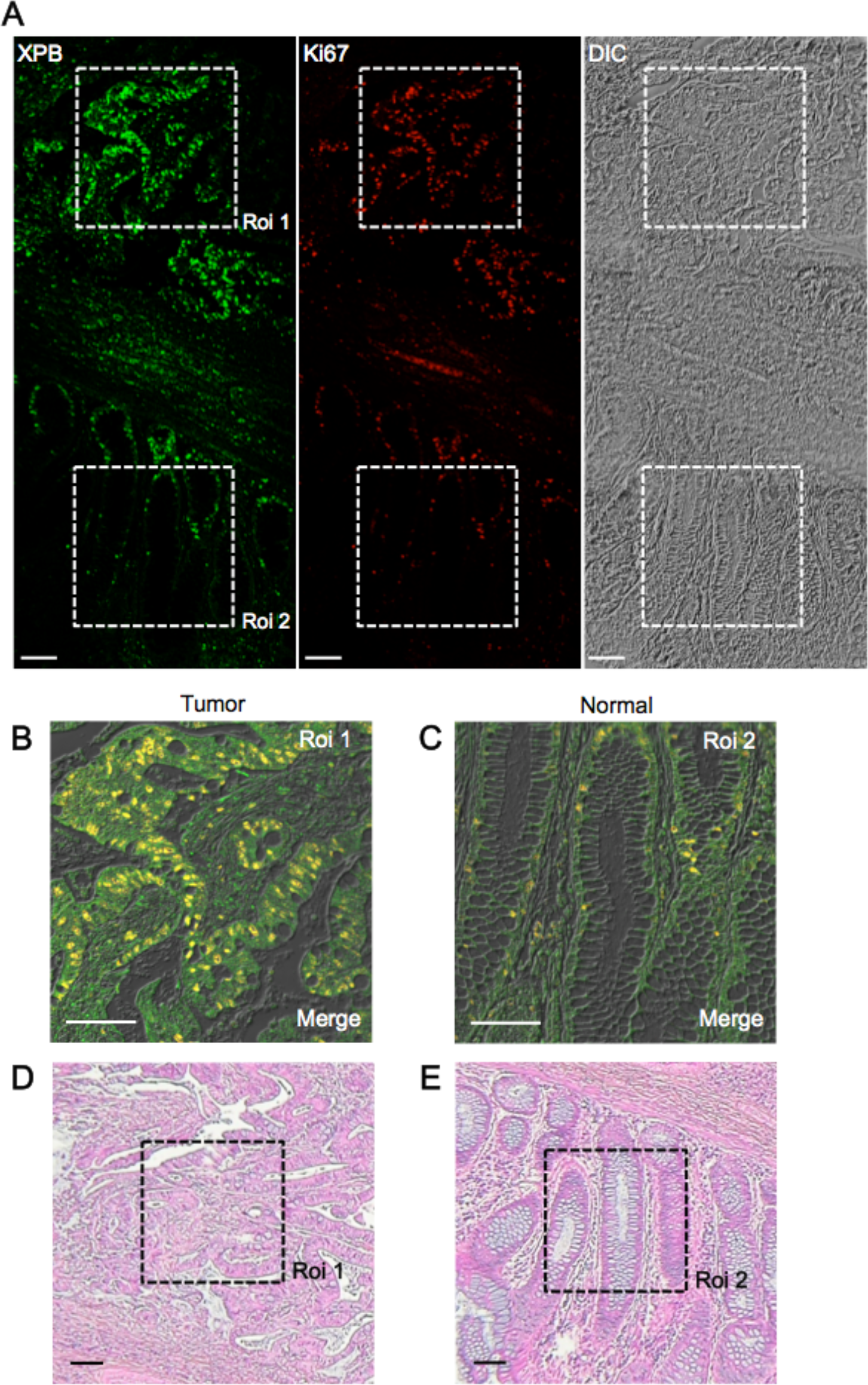
XPB and Ki67 immunoexpressions and pathologic features in human colon cancer specimens. **A.** Single tissue section presenting two distinct regions: a malignant region (ROI 1) and an adjacent normal colon mucosa (ROI 2). Elevated XPB and Ki67 expression appears frequently in the tumor cells of a colon cancer tissue (ROI1) compared to the cells of the adjacent normal mucosa (ROI 2). In the normal colon mucosa, a few cells showing concomitant high expression of XPB and Ki67 are seen in the deeper regions of colonic crypts where proliferating cells are expected to be present. **B, C.** Zoomed in ROI 1 and 2. Proliferating cells present high levels of both XPB (green) and Ki67 (red) and are easily identified by their yellow nuclei in both merged images. **D**, **E.** Consecutive serial section stained with hematoxylin to evaluate the morphology of the tissue. All scale bars represent 100 µm.

## Discussion

To maintain the chronic proliferative state, cancer cells need a high rate of active transcription by the three RNA polymerases. RNAP1 and RNAP2 have been found dysregulated in many cancer models. Enhanced RNAP1 activity triggers nucleoli enlargement and it is considered a marker of aggressive cancer cells associated with a poor prognosis (20). Enhanced RNAP2 activity has been found in many malignancies and many general transcription factors have been found consistently overexpressed in tumours (9). One striking example is the TBP protein, which has been found overexpressed in colon (21) and colorectal cancers (22). It has been proposed that overexpression of c-Myc, observed in many malignancies, could increase the proportion of elongating RNAP2 across the whole genome, in addition to stimulating particularly specific promoters (23). It seems nowadays widely accepted that a general increase in transcriptional activity is needed to maintain proliferation and survival of transformed cells.

In the present study, we have used a previously created knock-in mouse model expressing a fluorescently tagged basal transcription factor under its endogenous promoter, in order to disclose the quantitative relationship between steady-state levels of TFIIH and transcriptional activity of individual cells. By validating a method that combines such a knock-in mouse model, to 3D quantitative imaging techniques, we have been able to demonstrate that the concentration of TFIIH varies according to the metabolic state of different cell types of our model organism and that amongst cells of the same type, the total amount of TFIIH is tightly regulated to maintain a constant nuclear concentration. More importantly, we have revealed the existence of a direct relation between TFIIH concentrations, transcriptional activity and proliferation status.

Interestingly, we could find the TBP steady state is also proportional to the transcriptional activity of cells. However, the fact that TFIID (TBP is a subunit of this complex) composition changes depending on tissues and cellular differentiation state (24) might be problematic when wanting to find a general marker of transcriptional activity in a whole living organism. Surprisingly, RNAP2 steady state is not correlated with transcriptional activity. This result might be explained by the fact that the control on the activity of RNAP2 is operated by specific phosphorylations of the CTD (25), rather than a control on the cellular amount of RNAP2.

Many first-line chemotherapeutic drugs (i.e. cisplatin, actinomycin D) block both RNAP1 and RNAP2 transcription and their effectiveness is directly proportional to their cytotoxicity. In fact, blocking transcription induces apoptosis in proliferative cells but does not spare non-malignant cells, which may suffer from this kind of treatments. Specifically, two drugs have been developed to target TFIIH: (i) Triptolide that covalently binds to the XPB subunit of TFIIH and inhibits its ATPase activity (26) and triggering CDK7-dependent degradation (11) and hence disrupting both RNAP2 and RNAP1 transcription (27); (ii) Spironolactone that induce XPB degradation (12, 28). Nevertheless, these drugs have the same drawbacks that other drugs blocking transcription and their use may hinder the cellular activities of normal non-malignant cells. A safer and new approach to overcome cytotoxicity of chemotherapeutic drugs would be to modulate and reduce transcription instead of blocking it completely. In this context, we have hypothesised that low sub-optimal TFIIH concentrations at the cellular level may play a protective role by greatly hindering the unscheduled proliferation of cells required during carcinogenesis. Ensuing this hypothesis, we recently design two new drugs (compound 12 and 19) that interfere with the binding of the TTDA subunit to TFIIH (29). The rationale behind this study was that TTDA cells present a reduced level of TFIIH (30) and controls the stability of the TFIIH complex (2, 31). Reducing the steady state level of TFIIH by treating cells with these drugs decreased the level of TFIIH and transcriptional activity (29) and will in the future be tested in the context of carcinogenesis.

Surprisingly and remarkably, XPB concentration is tightly regulated in all cell types within tissues analysed in our study. Distinct cell types have distinct XPB concentrations and the concentration is kept strictly the same within the cell. This tight regulation might be explained by the fact that XPB, and hence TFIIH, by opening the helix of DNA during initiation and insuring the promoter escape of RNAP2 (32) might be one of the rate-limiting factor during the transcription process. The fact that cells within tissues have a rather fixed transcriptional program might explain why the concentration of certain transcription factors should be strictly controlled.

Finally, because of the tight relation between TFIIH steady state, transcriptional activity and proliferation, we propose to use our Xpb^y/y^ mouse model as a living biomarker of basal transcriptional activity and proliferation status. One of the major advantages of our mouse model for such studies is that it constitutes a unique source of biological material for a wide range of assays, from high throughput screening of small molecules, down to toxicological testing of candidate compounds directly with the Xpb^y/y^ mice.

## Methods

### Organotypic cultures

Organotypic explants of living tissues were produced as previously described(33, 34). Brain slices were analyzed on the same day of extraction in Neurobasal A (GIBCO) medium, supplemented with antibiotics and B27 at 37°C, 20% O_2_ and 5% CO_2_. Organotypic slices of spleen, kidney, liver and intestine, were produced by cutting 300 mm of the organ with a Tissue-chopper (McIlwain). Slices were analyzed within two hours following preparation in DMEM medium supplemented with 10% Fetal Calf Serum and 1% Penicillin/Streptomycin. Skin epidermis was prepared as described(35) and imaged in CnT07 medium (Bioconnect) supplemented with Ca^2+^ free 10% Fetal Calf Serum and 1% Penicillin/Streptomycin.

### Microscopy, image processing and analysis

All microscopy work was performed on a Zeiss 780 confocal, except for the data shown in Fig. 3b which was obtained on a Zeiss 710 NLO system (Zeiss, Germany). Both systems were used with a Plan-Neofluar 40×/1.30 oil immersion objective and had temperature and CO_2_ regulation. Briefly, image acquisition parameters were optimized for the (low) endogenous XPB-YFP signal, maintaining image quality as much as possible while keeping unwanted photobleaching to a minimum. Typical XYZ stacks of tissues cover a field of about 230×230 µm and 30 µm in the z direction to reduce the number of truncated nuclei during imaging (sampling intervals of 0.112 µm in x and y and 0.200 µm in Z; 2048×2048×150 pixels per stack). For cultured cells on glass, imaging was limited to between 10 and 13 µm. All 3D stacks were processed via quantitative deconvolution, performed using the classical maximum likelihood estimation algorithm provided by the Huygens Pro software (SVI, Hilversum, NL). Theoretical point spread functions (PSF) were used to allow the software to automatically generate different PSFs as a function of the imaging depth (“Brick mode” set to “More”). Signal to noise ratio was set to 10 for all channels and the quality change threshold kept at 0.1%. 3D reconstructions of the deconvolved image stacks were performed using the advanced object analyzer module of the same software. Typically, iso-surfaces for every cell nucleus are generated by selecting a threshold value corresponding to the background/non-specific fluorescence signal. Truncated and non-segmented nuclei are filtered out from the 3D reconstructions. Every volume enclosed by the remaining iso-surfaces is used to calculate the total fluorescence per cell nucleus (volume integral in arbitrary units) and to estimate the corresponding volume (in cubic micrometers). This volumetric quantification approach via 3D confocal microscopy and deconvolution was also validated by accurately measuring the doubling of DNA content during the cell cycle of a population of proliferating DAPI stained Chondrocytes (supplementary figure S1). For Fig. 3A, imaging and analysis of arrested and proliferative chondrocytes was simplified and greatly accelerated by approximating nuclear volumes to ellipsoids (approximate volume of a nucleus = 2/3 x largest transversal nuclear area x maximum nuclear thickness).

### Immunohistochemistry on murine tissues

Homozygous XPB-YFP mice and their corresponding controls were anesthetized and were perfused with 10% buffered neutral formalin solution. After perfusion, all organs were removed and post-fixed in formalin for 24 h. The organs were dehydrated and embedded in paraffin. Four-micron sections from paraffin blocks were deparaffinized in xylene and rehydrated through an alcohol series, washed in water and then subjected to microwave antigen retrieval at 98°C in citrate buffer (0.01 M in H2O, pH 6.0) for 45 minutes. Sections were immerged in 3% H2O2 for 5 minutes to block endogenous peroxidase activity and non-specific binding was blocked with Immunotech blocking serum for 5 minutes (universal HRP immunostaining, Immunotech, Beckmann Coulter, Marseille, France). Sections were rinsed in Phosphate buffered saline (PBS) and were incubated with the following primary antibody: mouse monoclonal anti-Green Fluorescent Protein overnight at 4°C (Clones 7.1 and 13.1; 1:50 dilution, Roche, Mannheim, Germany) and rat monoclonal anti-HA High Affinity for 60 minutes at room temperature (clone 3F10; 1:50 dilution, Roche). The sections were rinsed in PBS before incubation for 30 minutes with secondary Immunotech biotinylated antibody and 45 minutes with the Streptavidine-Peroxidase Complex (universal HRP immunostaining, Immunotech), followed by a 5 minutes-incubation with 3,3’-diaminobenzidine (DAB). Sections were washed in water, counterstained with Hematoxylin, dehydrated and mounted. Controls in each run included sections of wild type mice and sections incubated with PBS instead of primary antibody.

### Immunohistochemistry on human tissues

Anti-TFIIH immunohistochemistry was performed on formalin-fixed paraffin embedded tissue microarrays of various tumors. Four micrometer sections from paraffin blocks of were deparaffinized in xylene and rehydrated through an alcohol series, washed in water and then subjected to microwave antigen retrieval at 98°C in citrate buffer (0.01 M in H_2_O, pH 6.0) for 45 minutes. Sections were immerged in 3% H_2_O_2_ for 5 minutes to block endogenous peroxidase activity and non-specific binding was blocked with diluted normal horse serum for 20 minutes (Vectastain Universal Elite ABC kit, Vector Laboratories, Burlingame, California, USA). Sections were incubated 1 hour at room temperature with the following primary antibody: rabbit polyclonal anti-TFIIH p89 (1:50 dilution, Santa Cruz Biotechnology, Inc. Santa Cruz, California, USA). The sections were rinsed in PBS before incubation for 30 minutes with diluted biotinylated “universal” secondary antibody and 30 minutes with the ready-to-use Vectastain Elite ABC Reagent (Vector Laboratories), followed by a 5 minutes-incubation with 3,3’-diaminobenzidine (DAB). Sections were washed in water, counterstained with Hematoxylin, dehydrated and mounted.

### Immunofluorescence anti-EU in MDF

XPB-YFP murines Mouse Dermal Fibroblasts (MDF) were grown for 24h on coverslips before being incubated for 30 min with 100µM of 5-ethynyl uridine (EU from ThermoFisher). After washes with PBS, cells were fixed with 3.7% paraformaldehyde for 15 min at 37°C, then incubated 2 times for 5 min with PBS and 3% BSA. Cells are then permeabilized for 20 min with PBS 0.5% triton X-100. After 30 min of incubation with PBS 0.5% BSA and 0.15% glycine, cells were in contact for 2 hours at room temperature with a primary antibody: Pol2 (8WG16, 1/100 mouse, santacruz sc56767), HA for XPB staining (1/200, rat, Roche 3F10), TBP (1/200, rabbit, cell signalling 8515S) and p62 (1/100, rabbit, abcam ab204168). After several washes with PBS 0.1% triton X-100, cells were incubated for 1h with a secondary antibody: Alexa-488nm donkey anti-rat or goat anti-mouse or goat anti-rabbit (Invitrogen) (dilution 1/400). Cells were washed several times with PBS 0.1% triton X-100 and then incubated for 30min with Click-iT reaction cocktail containing Alexa Fluor azide 594. After washing, the coverslips were mounted with Vectashield (Vector laboratories).

### Immunofluorescence anti-BrU in Chondrocytes

XPB-YFP murines chondrocytes were grown for 48h on coverslips before being incubated for 30 min with 2.5 mM bromo-uridine (Sigma-Aldrich). After washes with PBS, cells were fixed with 2% paraformaldehyde 15 min at room temperature, then washed 3 times rapidly and 2 times for 10 min with PBS 0.1% triton X-100. After 30 min of incubation with PBS 0.5% BSA and 0.15% glycine, cells were in contact for 2 hours at room temperature with a mouse monoclonal primary antibody anti-BrdU (1/1000 dilution) (Roche), then rinsed and incubated for 30 min with a fluorescent secondary antibody Alexa-633nm goat anti-mouse (Invitrogen) (dilution 1/400), and finally mounted with Vectashield (Vector laboratories).

### Protein extraction

For protein extraction, cells were cultured in 10-cm dishes. Cells were harvested by scratching. For arrested vs proliferation state, cells were counted and the same number of cells were lysis. The extraction of total proteins has been performed using the ProteoJET mammalian Cell Lysis Reagent (Fermentas). For WT vs XPB-YFP, the concentrations of proteins were determined by the Bradford method. The samples were then diluted with Laemmli buffer (10% glycerol, 5% β-mercaptoethanol, 3% sodium dodecyl sulfate, 100mM Tris-HCl [pH 6.8], bromophenol blue), heated to 95°C, and loaded on an SDS-PAGE gel.

### SDS-PAGE

Proteins were separated using SDS-PAGE gel composed of bisacrylamide (37:5:1) and blotted onto a polyvinylidene difluoride membrane (0.45-µm pore size; Millipore). Nonspecific sites were blocked in skimmed milk in the presence of 0.1% Tween 20, and the membrane was incubated with an appropriate antibody. We used the following antibodies: XPB p89 (1/500 rabbit, santacruz sc293), CDK7 (1/2000, rabbit, santacruz sc856X), CyclinH (1/1000, mouse, IGBMC 2D4) and p62 (1/500, rabbit, abcam ab204168). The loading was controlled with anti-α-tubulin antibody (1/50000, mouse, sigma T6074). Protein bands were visualized via enhanced chemiluminescence (Pierce ECL Western blotting substrate) using horseradish peroxidase-conjugated secondary antibodies and imaged using the ChemiDoc system (Bio-Rad). The quantification of the band was performed with ImageLab software (Bio-Rad) using the method of volumes (rectangle). The background was removed using the local subtraction method.

## Acknowledgments

This study was supported by the Institut National du Cancer (*PLBIO17-043*), l’Agence Nationale de la Recherche (ANR DyReCT: ANR-14-CE10-0009) and the ARC foundation (Association pour la Recherche sur le Cancer, “projet Fondation ARC PJA 20131200188).

## Author Contributions

GGM, POM designed the experiments. LMD, POM, CM performed the experiments and POM analysed the data. WV provided with some reagents. GGM and POM wrote the paper.

## SUPPLEMENTARY MATERIAL

**Figure S1.**
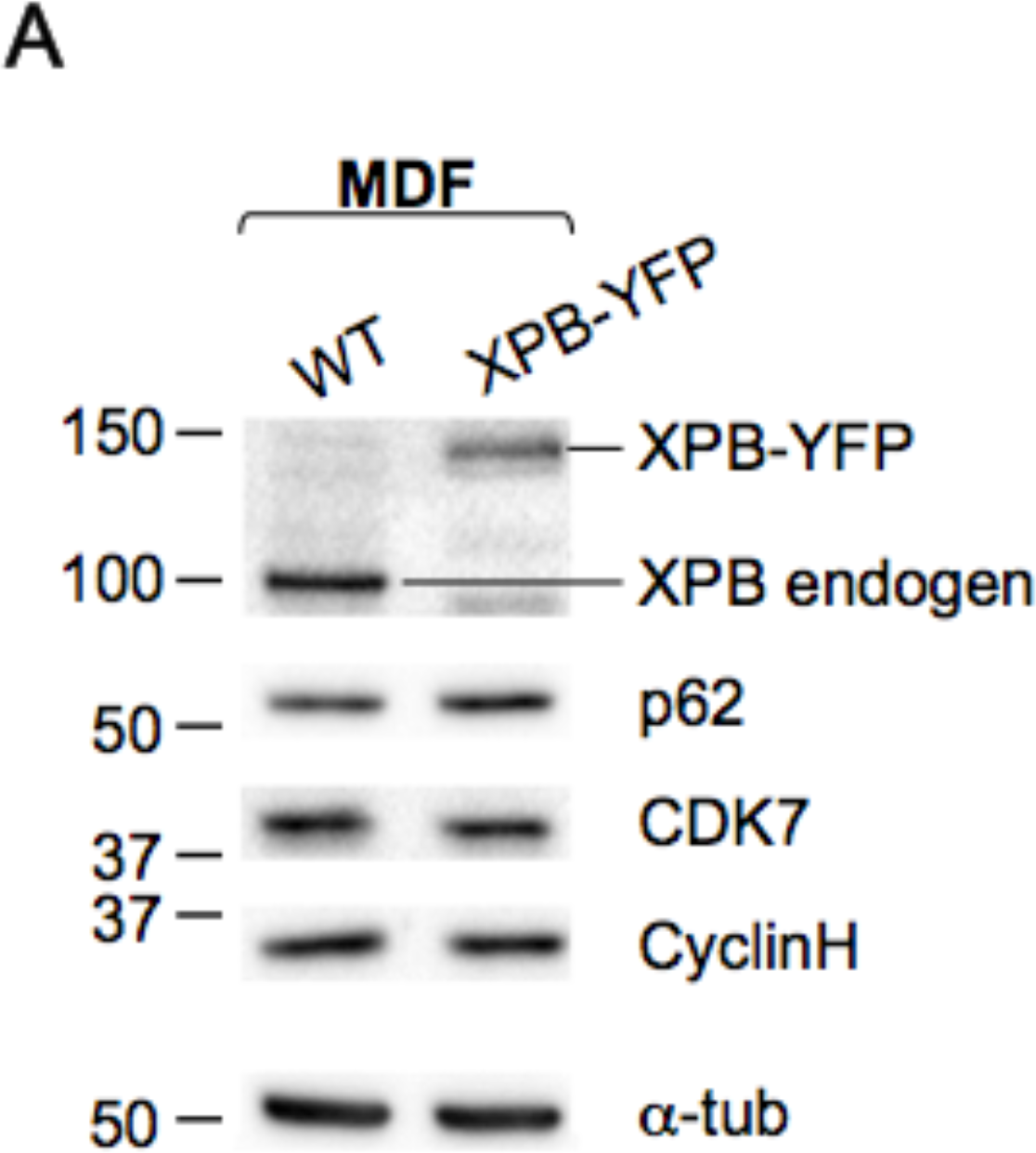
Western blot showing the expression of TFIIH subunits in whole cell extract of WT and XPB-YFP stably-expressing MDF cells. α-tub serves as a loading control.

**Figure S2.**
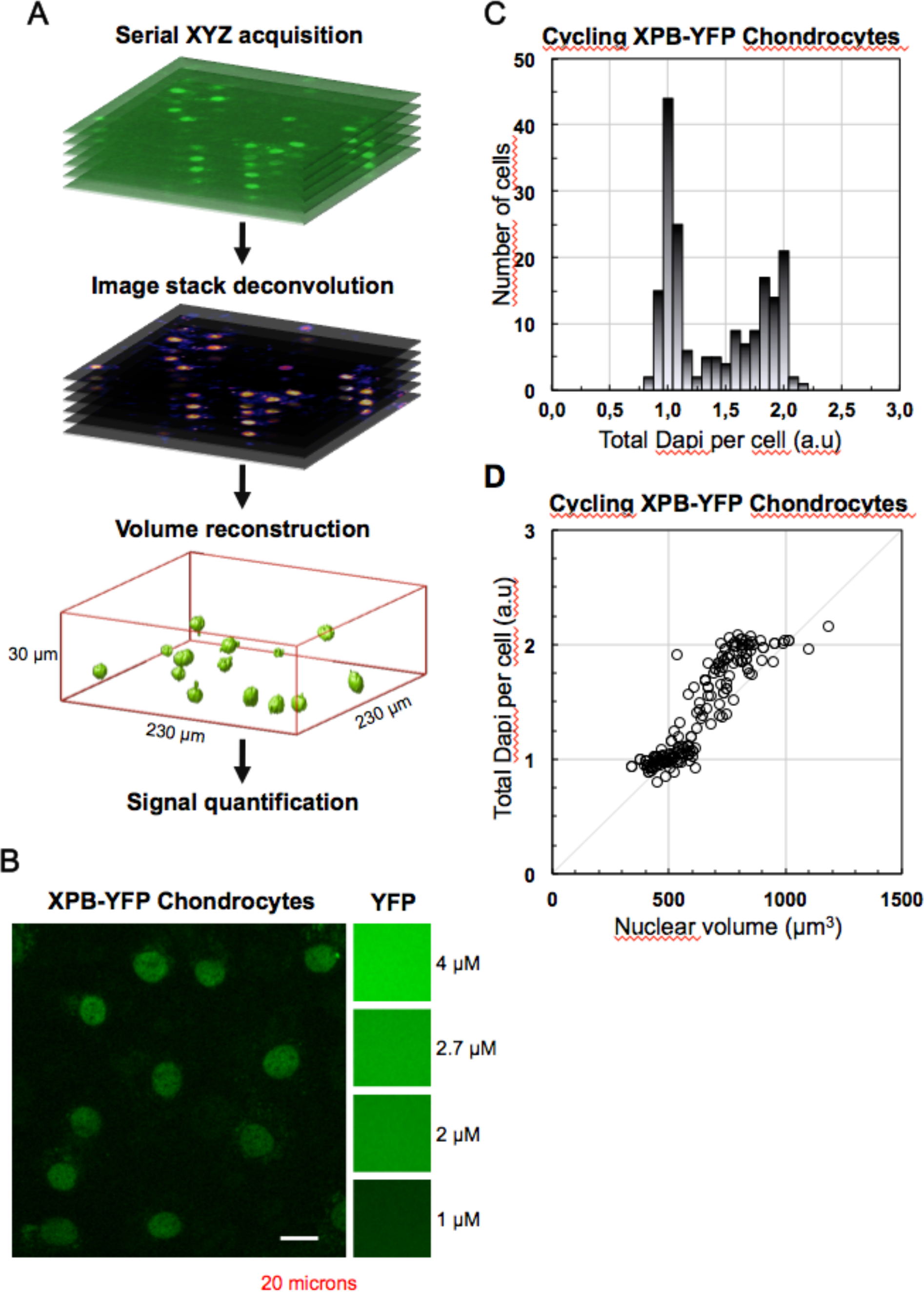
**A** Simplified scheme representing the image post-processing and analysis pipeline. First, living organotypic tissue slices or cultured cells (Xpb^y/y^ mouse) are imaged by confocal microscopy, typically over a 230×230×30 µm region. Second, the resulting image stacks are processed via deconvolution to restore the spatial distribution of the fluorescence signal. Third, using the same software, fluorescence intensity iso-surfaces are generated to calculate the volume occupied by the XPB-YFP signal of well segmented, non-truncated cell nuclei. Finally, the total XPB-YFP derived fluorescence per cell was estimated by integrating over each of these volumes. **B**. Side by side comparison of the XPB-YFP signal in living chondrocytes (Xpb^y/y^ mouse) and the fluorescence of diluted YFP recombinant protein obtained with identical imaging parameters. Scale bar represents 20 µm. **C.** Single cell fluorescence quantification of DAPI stained cycling murine chondrocytes via deconvolved 3D imaging as described in the first panel. The graph is a histogram of the resulting total DAPI signal per cell (in arbitrary units). After normalizing the lowest peak to 1, the second peak is found at 2±0.05. **D.** Scatter plot of the normalized total DAPI signal per cell versus the reconstructed nuclear volume. The dotted line represents a constant DAPI concentration.

